# The evolutionary origins of the lysosome-related organelle sorting machinery reveal fundamental homology in post-endosome trafficking pathways

**DOI:** 10.1101/2024.01.30.578091

**Authors:** Kiran J. More, Joel B. Dacks, Paul T. Manna

## Abstract

The major organelles and pathways of the endomembrane system were in place by the time of the last eukaryotic common ancestor (LECA) (∼1.5 billion years ago) and their acquisition were defining milestones during the process of eukaryogenesis itself. Comparative cell biology and evolutionary analyses show multiple instances of homology in the protein machinery controlling distinct inter-organelle trafficking routes. Resolving these homologous relationships allows us to explore processes underlying the emergence of new cellular compartments, infer ancestral states pre-dating LECA, and can even provide insight into the process of eukaryogenesis itself. Here we undertake a molecular evolutionary analysis, including providing a transcriptome of the jakobid flagellate *Reclinomonas americana,* exploring the origins of the machinery responsible for the biogenesis of lysosome-related organelles, the so-called Biogenesis of Lysosome-related Organelle Complexes (BLOCs 1,2, and 3). This pathway has been studied only in animals and is not considered a feature of the basic eukaryotic cell plan. We show that this machinery, and by inference the corresponding sorting pathway, was likely in place prior to the divergence of eukaryotes and is found in a much more diverse array of eukaryotes than is currently assumed. As such, this sorting pathway is likely an underappreciated facet of broader eukaryotic cellular function. Moreover, we resolve multiple points of ancient homology between all three BLOCs and other post-endosomal retrograde trafficking machinery (BORC, CCZ1/MON1, and a newly identified relationship with HOPS/CORVET) offering a mechanistic and evolutionary unification of these trafficking pathways. Overall, this study provides a comprehensive account of the rise of the LRO biogenesis machinery from prokaryotic origins to current eukaryotic diversity, Asgard archaea to animals, integrating it into the larger mechanistic framework describing endomembrane evolution.

## Introduction

Sub-cellular compartmentalisation is a defining feature of eukaryotes. The basic sub-cellular plan of the eukaryotic cell (e.g. the Golgi apparatus, endosomal system, nucleus, ER) is well conserved and was fully established by the time of the last eukaryotic common ancestor (LECA), prior to the divergence of the extant eukaryotic lineages (Koumandou et al., 2013). The evolution of each of these major endomembrane organelles are key milestones that must be explained in any scenario of eukaryogenesis. The molecular machinery responsible for the biogenesis and maintenance of much of this compartmentalisation is now relatively well understood. Most of this machinery has arisen through paralogous expansion of a limited number of gene families, with individual paralogs operating at distinct locations within the cell. These core membrane trafficking gene families for which the evolutionary paths have been examined include the hetero-tetrameric adaptor protein complexes (HTACs) (Hirst et al., 2011) (Hirst et al., 2014) (De Franceschi et al., 2014) (Schlacht and Dacks, 2015), SNAREs (Kloepper et al., 2007), Ras family GTPases (van dam et al., 2011) (Elias et al., 2012) (Klöpper et al., 2012), and the CATCHR families of multi-subunit tethering complexes (MTCs) (Koumandou et al., 2007) (Santana-Molina et al., 2021).

The Organelle Paralogy Hypothesis (OPH) integrates these observations and suggests that the emergence of novel endomembrane sorting pathways—and as a result, new compartments—is the result of paralogous expansions of gene families and neofunctionalization of redundant paralogues (Dacks and Field, 2007). Data supporting this as the mode of endomembrane organelle emergence can be found from phylogenetics, but also from computational modelling, and evolutionary cell biology of lineage-specific organelles in plants and apicomplexan parasites (Ramadas and Thattai, 2013) (Hirst et al., 2014) (Agop-Nersesian et al., 2010) (Kanazawa et al., 2020) (Koreny et al., 2023) (Klinger et al., 2022). The OPH offers a mechanistic explanation for the elaboration of the non-endosymbiotically derived eukaryotic endomembrane system organelles. Phylogenetic analyses of Rabs (Elias et al., 2012), Arfs (Vargová et al., 2021), and HTAC proteins (De Franceschi et al., 2014) (Hirst et al., 2014) have allowed for some resolution moving from the LECA back through eukaryogenesis, reconstructing some of the more proximal events. The identification of proteins or domains in the Asgard archaea, the best candidate for the archaeal lineage from which the eukaryotes are derived, that are clearly homologous to many components in the molecular machinery of membrane-trafficking anchors the origins of the membrane-trafficking machinery on each end of eukaryogenesis, but leaves tremendous gaps in the detail in between the FECA and LECA (Zaremba-Niedzwiedzka et al., 2017) (Hatano et al., 2022).

Moreover, there are some sets of membrane trafficking system machinery that have not yet been integrated into this framework. One of the most glaring omissions are the Biogenesis of Lysosome-related Organelle Complexes, or BLOCs. These proteins were first identified through tracing mutations found in a cluster of similar genetic diseases, called Hermansky-Pudlak Syndrome, that result in impairment of trafficking to a selection of organelles termed Lysosome-related organelles (LROs) (Bowman et al., 2019). Identified and characterised in animals, LROs are a diverse array of secretory organelles that are derived from the endosomal and lysosomal sorting pathways. In humans, LROs include the dense granules of platelets, lytic granules of T-Lymphocytes, melanosomes of melanocytes, and Weibel-Palade bodies of endothelial cells (Bowman et al., 2019).

The LRO-specific sorting machinery comprises three unrelated multi-subunit protein complexes: BLOC-1, BLOC-2, and BLOC-3. BLOC-1 is a hetero-octameric complex involved in cargo sorting and formation of tubular transport carriers at the early endosome (Falcón-Pérez et al., 2002) (Ciciotte et al., 2003) (Li et al., 2003) (Starcevic and Dell’Angelica, 2004) (Bowman et al., 2021) (Jani et al., 2022). BLOC-2 is a trimeric complex that acts downstream of BLOC-1 and is suggested to perform a tethering function between BLOC-1 generated tubules and acceptor LROs (Gautam et al., 2004) (Di Pietro et al., 2004) (Dennis et al., 2015). BLOC-3 is a hetero-dimer that acts as a GEF for Rabs 32A and 38 (Gerondopoulos et al., 2012), which are also required for LRO biogenesis (Wasmeier et al., 2006). BLOC-3 subunits have homology to the nominal two subunits of the trimeric MON1-CCZ1 complex (Cheli and Dell’Angelica, 2010), which acts as a Rab GEF for Rab7 during trafficking from the late endosome to the lysosome. This homology suggests that the LRO pathway may have arisen via a mechanism in line with the organelle paralogy hypothesis. The current understanding of the evolutionary origins of the other BLOCs is insufficient to address this question fully.

The timing of the origin of the BLOCs is also unclear. From earlier studies, it is clear that the majority of the protein subunits of BLOC-1, -2, and -3 were present at the origins of Amorphea (the evolutionary lineage encompassing amoebozoans, fungi and animals) (Cheli and Dell’Angelica, 2010) (Pu et al., 2015). Beyond this lineage, the presence of the BLOCs has not been systematically assessed. Three subunits of BLOC-1 were identified in organisms outside of Amorphea (Cheli and Dell’Angelica, 2010); however, the later identification of a protein complex that shares these three subunits with BLOC-1, named the BLOC-one-related complex (BORC) (Pu et al., 2015), means that we do not currently have any evidence of BLOC-1 specific subunits outside of the Amorphea. In contrast, at least two BORC specific subunits were identified in diverse eukaryotic taxa (Pu et al., 2015), suggesting that this complex pre-dates the diversification of the eukaryotes. Whilst BLOC-1 and BORC share three subunits and their respective components display similar predicted structure, the precise relationships between BLOC-1 and BORC subunits have not been explored. In addition to these BLOC-1/BORC subunits, potential orthologs of the BLOC-3 subunits HPS1 and HPS4 have been identified in the metamonad *Trichomonas vaginalis* and the discobid *Naegleria gruberi* respectively (Cheli and Dell’Angelica, 2010) (De Franceschi et al., 2014). The discovery of these subunits in two deep-branching eukaryotic lineages indicates a potential for these complexes to be ancient and paneukaryotic, but a single subunit is not enough to confirm a pre-LECA origin for the complex.

There is currently no clear picture of the extent to which the BLOCs, BORC, and associated machinery are conserved components of the endomembrane machinery across eukaryotes, nor as to how these complexes fit in the larger picture of endomembrane system evolution. We have therefore undertaken a detailed evolutionary analysis of the full complement of BLOC subunits across a broadly representative sample of eukaryotes, taking advantage of the vast number of genomes and transcriptomes that have been sequenced in the years since the initial surveys were performed. This included producing a new transcriptome for the jakobid *Reclinomonas americana* which represented a crucial missing sampling point in our analysis. Finally, we have explored the deep evolutionary relationships between the BLOCs and other membrane trafficking machinery in order to integrate them into the larger picture of membrane-trafficking system evolution during eukaryogenesis.

## Methods

### Comparative genomics

Genomes and transcriptomes were collected from a wide variety of publicly and privately available resources (S1 Table), including an in-house assembly of a transcriptome of *Reclinomonas americana* (methods in S1 Text). Searches were performed in three parts. Initial BLAST searches were performed with the human proteins as starting queries into a reduced set of predicted proteomes. Positive hits were confirmed with reciprocal best hit (RBH) against the human proteome. The middle stage was iterative searching using HMMer and/or lineage-specific BLAST searches until no new positive hits could be identified in the reduced set of proteomes. Finally, human proteins and the constructed HMMs (S1 Dataset) were queried into the full genome selection. Reverse searches were done into a custom database containing diverse organisms with proteins annotated during the initial search step. In the case of BLOC-1 and BORC subunits, JackHMMER (Johnson et al., 2010) was used for reverse searches against UniProt eukaryotic reference proteomes (release 2023_03). Hits were considered positive if the reverse search yielded the human protein within three iterations, and had at least two orders of magnitude e-value difference between the next non-redundant hit. BLAST and HMMER results were performed and parsed using the AMOEBAE workflow using a maximum e-value of 0.05 and a minimum difference in e-value order of magnitude of 2 (Barlow et al., 2023), while JackHMMER results were parsed using in-house scripts (available on Figshare). Phylogenies were used to confirm identities between known paralogs (HPS1/MON1, HPS4/CCZ1, and HPS5/TECPR2). Positive hits were aligned using mafft-linsi with 1000 iterations; fragments were aligned using –addfragments (Katoh and Standley, 2013). Alignments were trimmed using an in-house script that retained sites with 50% identity and fewer than 30% gaps, and then adjusted manually. Maximum likelihood trees were inferred with 1000 ultrafast bootstraps in IQ-TREE 2 under the best performing model inferred by ModelFinder using BIC (Kalyaanamoorthy et al., 2017).

### Characterization of BLOC-2

Positive hits of the BLOC-2 subunits were further characterized for secondary structure using Ali2D (Gabler et al., 2020), and domain structure using InterProScan (Jones et al., 2014). Representative pan-eukaryotic HMMs of the BLOC-2 subunits were queried using the HHblits webserver (Remmert et al., 2011) (Gabler et al., 2020) using 8 iterations and retaining 10000 hits under the default inclusion parameters.

### Characterization of BLOC-1 and BORC

A database of all identified BLOC-1 and BORC subunits was generated for use in all-versus-all JackHMMER searches. The searches were run for three iterations, with an e-value cut off of 1 x 10^-3^. Markov Clustering (MCL) was then applied using -log transformed E-values as edge weights. The MCL algorithm was implemented via the clusterMaker Cytoscape plugin (Morris et al., 2011).

## Results

### BLOC-1 and BORC are pan-eukaryotic

To begin investigating the evolution and conservation of the BLOC-1,- 2, -3, and BORC subunits, specifically their prevalence across eukaryotes, we carried out homology searches using an iterative HMM-based approach against a panel of broadly representative eukaryotic predicted proteomes. We identified over 1000 putative homologous sequences from over 140 taxa, greatly expanding the previously reported survey of these complexes (**S2 Table, S3 Table**). Following sequence validation, the phyletic distribution of identified homologues was assessed in order to infer the likely points of origin of the various subunits and complexes.

The shared subunits of BLOC-1 and BORC were most widely identified, with representatives of all three subunits present in every supergroup. However, even these shared subunits showed incidences of inferred secondary loss within some groups, particularly within Diaphoretickes, including rhodophyte and phaeophyte algae, fungi, and apicomplexan parasites (**Figure 1**).

**Figure 1.**
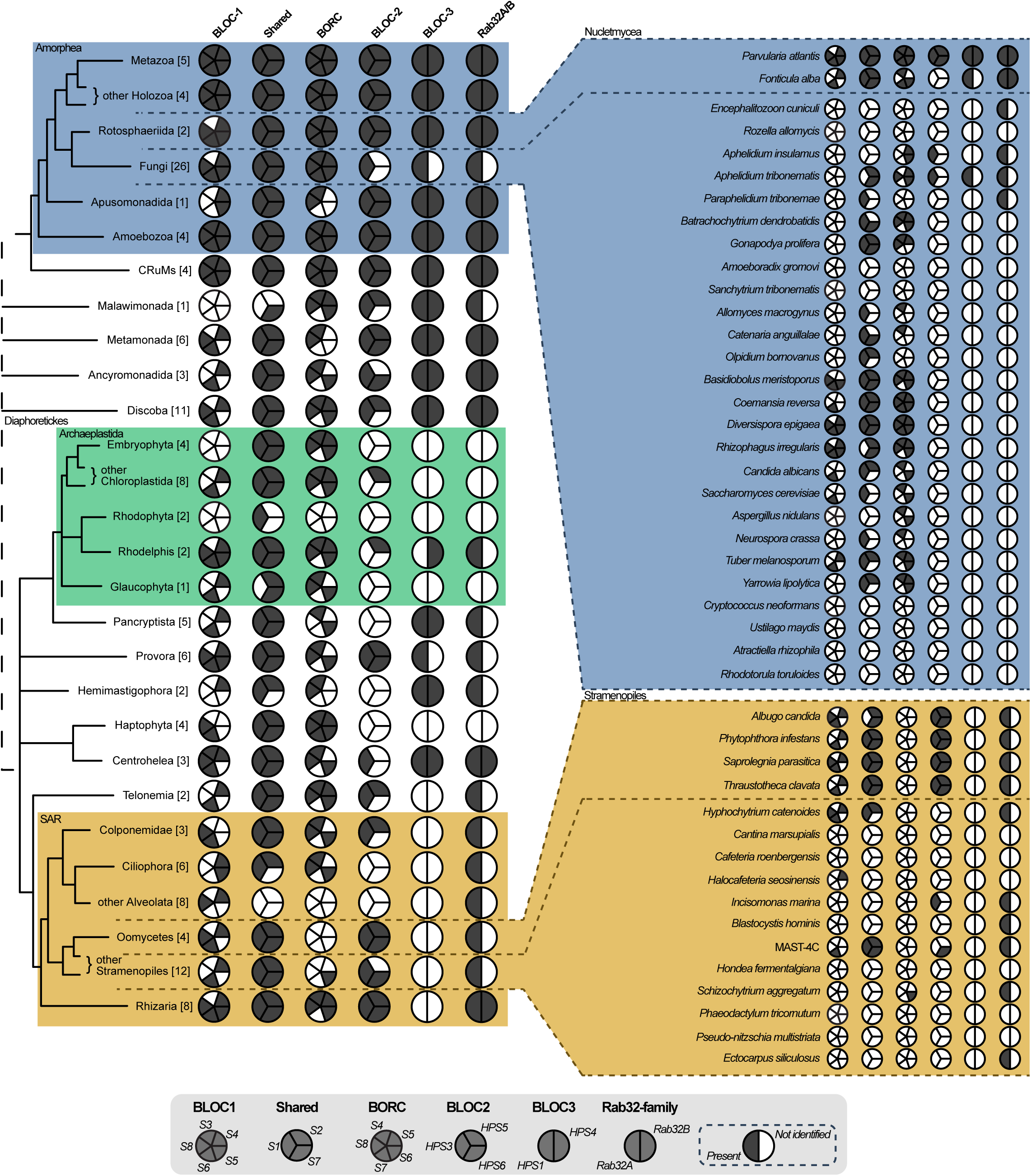
Distribution of BLOC subunits and the LRO-associated Rab32 family. Filled segments denote that the corresponding subunit has been identified with high confidence in at least one taxon within the indicated group. Clades of particular interest for the current study are highlighted in colour. Gene complements in individual taxa from selected collapsed groupings are expanded to the right. Full data at the level of individual taxa can be found in S3 Table.

At least some representation of BORC specific subunits was found in all eukaryotic supergroups, with repeated incidences of loss amongst the taxa within Diaphoretickes (Figure 1; S3 Table). Thus, BORC likely arose pre-LECA. BORCS7 was the only subunit to have arisen later, with its identification almost completely restricted to the superbranch containing Amorphea and CRuMs (**Figure 1**). Given only a single other hit was found outside this distribution (in Haptophyta), this is likely a false positive.

Similarly, we identified representatives of all BLOC-1 specific subunits beyond the Amorphea, with all but BLOC1S3 and BLOC1S5 broadly distributed enough to confidently support a pre-LECA origin. Only BLOC1S3 was almost completely restricted to Holozoa, with merely three hits identified in the remaining 138 eukaryotes searched, likely indicating that these are false positives. BLOC1S5 was identified in all major lineages of Amorphea, as well as the related group CRuMs, but was found only sparsely in Diaphoretickes. Since these hits were never identified in closely related taxa, we are only able to confidently infer an origin for BLOC1S5 at the divergence of Amorphea and CRuMs. Large scale losses of BLOC-1 were seen amongst the Diaphoretickes (including in land plants), but not necessarily co-incident with the loss of BORC. We rarely observed BLOC-1 or BORC specific subunits in species that totally lacked the shared subunits, increasing our confidence in our identifications.

Despite our best efforts, we should note that the possibility of both false negatives and false positives remain, particularly as these proteins are small and display relatively low sequence conservation. By focusing our analysis and interpretation on patterns of presence/absence rather than giving large amounts of weight to any single protein, we are nonetheless confident that we can infer that BLOC and BORC components are widespread amongst eukaryotes and were likely present in LECA.

### BLOC-2 and -3 are also widely distributed in eukaryotes and inferred to have been in LECA

As with BLOC-1, our BLOC-2 and -3 analysis extends the existing taxonomic sampling substantially. Our initial BLAST-based searches suggested that these complexes were largely found within Amorphea and CRuMs, which have been inferred to be on the same side of the eukaryotic root (Derelle et al., 2015), and generally absent from Diaphoretickes. A remarkable exception to this was the clear but taxonomically restricted identification of HPS3 and HPS5 in oomycetes. Placement of the point of origin for these complexes was therefore impossible using BLAST-based searches alone. HMM-based searches proved far more sensitive in the detection of BLOC-2 and BLOC-3 subunits, detecting some representation of both complexes spread widely across the sampled taxa (**Figure 1**). This included the BLOC-2 subunit HPS6 as well, which had previously only been identified in vertebrates (Cheli and Dell’Angelica, 2010) (S2 Text).

Despite all BLOC-2 subunits being found in oomycetes in our initial searches, there was an almost total lack of other BLOC-2 and -3 subunits within Diaphoretickes. There were, however, BLOC-2 and BLOC-3 proteins identified in Discoba, which is a deep-branching lineage of mostly free-living eukaryotes that is associated with Diaphoretickes (Derelle et al., 2015). Overall, BLOC-2 and -3 subunits were however very sparsely identified, with “complete” versions of both complexes almost exclusively found in Amorphea, apart from BLOC-2 in oomycetes and BLOC-3 in metamonads and discobids, suggesting an unusually complex pattern of retention and recent losses. To further examine this unusual distribution, we made every attempt to expand the taxon selection within Diaphoretickes and Discoba using available genome and transcriptome resources. There is currently only a single genome representing Jakobida, the deepest branching lineage within Discoba. Therefore, we produced our own transcriptome of the jakobid *Reclinomonas americana,* containing 21,101 predicted proteins after deduplication. The BUSCO score (90% complete, 4% fragmented, 6% missing against OrthoDB eukaryote version 10), compared very favourably to that of the highly contiguous genome assembly of the jakobid *Andalucia godoyi* (88.8% complete in OrthoDB 9.1) (Gray et al., 2020). Searching this transcriptome yielded further identifiable BLOC-2 and -3 subunits including both HPS1 and HPS4 subunits of BLOC-3. As in the case of BLOC-1 and BORC, the broad distribution of BLOC-2 and BLOC-3 subunits supports a pre-LECA origin for these complexes.

For BLOC-3, this assertion is further supported by phylogenetic reconstruction of the HPS1/MON1 and HPS4/CCZ1 families (**S1 Figure**). BLOC-3 subunits HPS1 and HPS4 have previously been proposed to be paralogs of MON1 and CCZ1, respectively (Hoffman-Sommer et al. 2005; Kinch and Grishin 2006; Cheli and Dell’Angelica 2010). Phylogenetics with our diverse set of eukaryotes confirm this as an ancient split between the two complexes, which diverged prior to the LECA (**S1 Figure**). We also note that in all cases our reconstructed BLOC-2 subunit phylogenies broadly reflect the eukaryotic tree, arguing against horizontal gene transfer as a driver of the observed phyletic distribution and further supporting a pre-LECA origin for this complex **(S2 Figure).**

Finally, we updated the established distribution of the LRO biogenesis associated Rab32 family with our expanded taxon sampling. This Rab family is known to have emerged pre-LECA, and in humans is represented by the Rab32A paralogs Rab29, Rab32, and Rab38 (Elias et al., 2012) (Ortiz-Sandoval et al., 2014). Through RBH BLAST searches, we found that its retention across the eukaryotic tree was for the most part in good agreement with our observations of the BLOCs, as well as past literature results (Figure 1, S3 Table) (Elias et al., 2012) (Ortiz-Sandoval et al., 2014).

In summary, our data suggest that all three BLOCs arose prior to the diversification of eukaryotes and are broadly but patchily retained across extant eukaryotes.

### Reconstructing deep inter-subunit relationships between BLOC-1 and BORC

In order to better understand the LRO-associated biogenesis pathway in the greater context of endomembrane system evolution, we investigated the deeper homology of the complexes to those of other pathways.

To begin, some homology between components of the BLOCs and BORC has been suggested. When the BORC complex was identified it was noted that the BORC5 and BORC6 subunits retrieved putative yeast BLOC-1 subunits (Snn1p and Cnl1p respectively) by HHM based homology searches (Pu et al., 2015). However, no comprehensive analysis of the relationships between subunits of the two complexes has been carried out. Additionally, our data suggest that the yeast sequences are likely mis-attributed and are in fact BORC subunits (**Figure 1**, S2 Table, S3 Figure). We therefore attempted to identify any relationships between BLOC-1 and BORC subunits using our comprehensive, well validated, and broadly representative dataset of sequences.

Our initial BLAST searches had indicated some clear sequence-level relationships between subunits. The shared subunit BLOC1S7 appeared closely related to the BLOC-1 specific BLOC1S6 across multiple lineages. We also detected remote similarity between BLOC1S4 and BORCS6, suggesting additional homology. We attempted to reconstruct these relationships phylogenetically, however this approach was hampered by relatively short sequence length and poor sequence conservation. We therefore adopted an alternate HMM-search and sequence clustering approach to identify distant relationships between BLOC-1 and BORC subunits.

We generated a database of all identified BLOC-1 and BORC subunits and carried out an iterative all vs. all JackHMMER search within this database. After 3 iterations, suggestive patterns of sequence similarity were apparent (**Figure 2A).** Further iterations did not add any resolution to the data. The majority of BLOC-1 and BORC subunits showed a degree of similarity to at least one other subunit. The exceptions are the likely Amorphea-specific subunits BLOC1S3, BLOC1S5, and BORCS7, as well as the apparently ancient BORCS8 subunit. To further interrogate these relationships in an unbiased manner we carried out a -log(E-value) edge weighted Markov Clustering (Enright et al., 2002) of the dataset (Figure 2B). This suggested two major groupings of similar sequences, the first comprising BLOC1S6 and BLOC1S7, as mentioned above, alongside the BORC specific subunit BORCS4 (KXD1 in humans). The second group comprised the BLOC-1-specific BLOC1S4 and the BORC specific BORCS6 (as observed in our initial similarity searches), as well as the shared subunit BLOC1S2. In both cases these groups contain one shared, one BLOC-1- and one BORC-specific subunit. This is the first evidence of paralogy of these complexes extending beyond the shared subunits of BLOC-1 and BORC, revealing the parallel expansion of the pathways from a primordial lysosome-associated complex.

**Figure 2.**
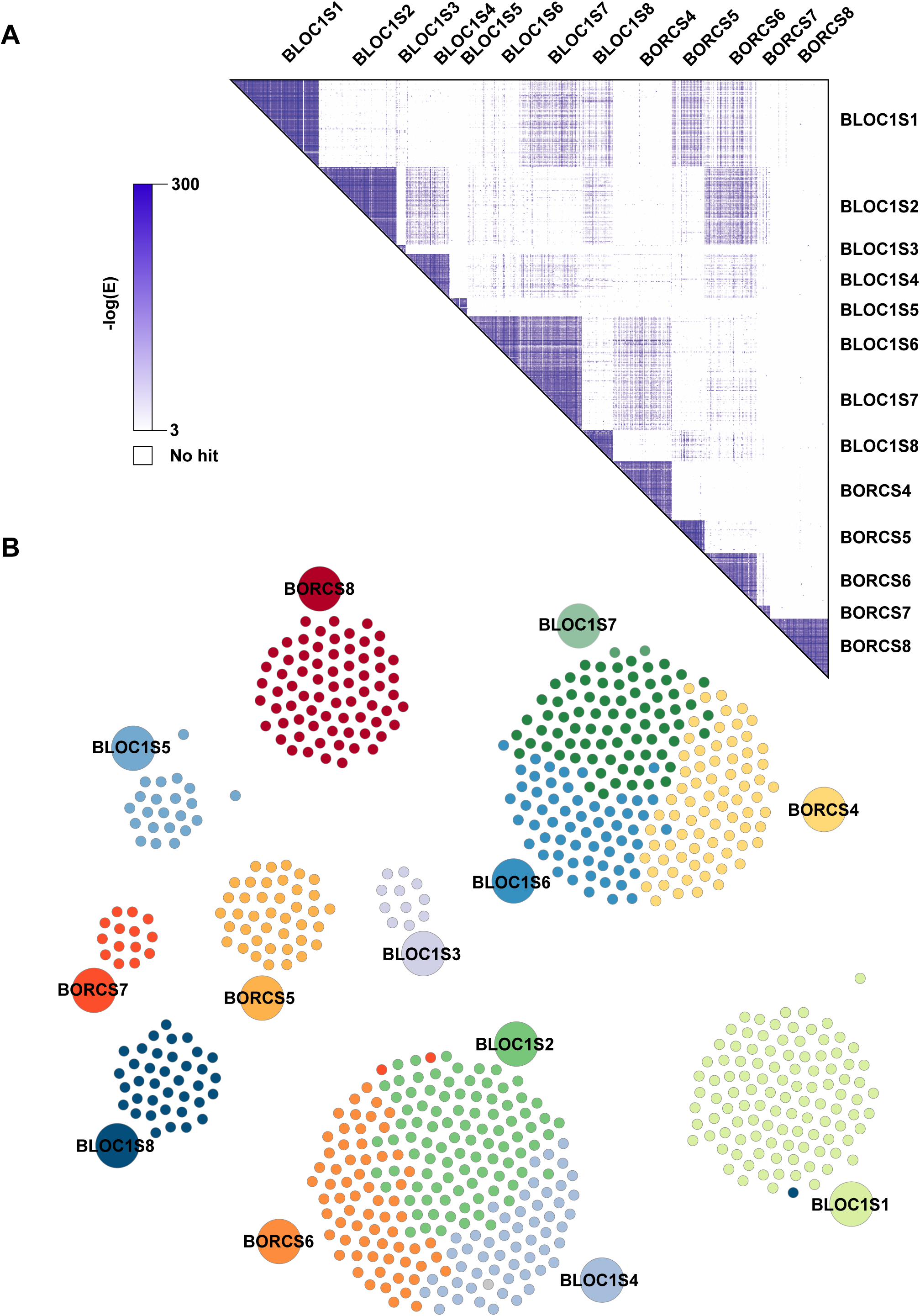
Inter-subunit similarity of BLOC-1 and BORC. **(A)** Heatmap showing -log(E-value) scores after 3 iterations of all vs. all JackHMMER searches. **(B)** -log(E-value) edge weighted Markov Clustering (MCL) of our identified BLOC-1 and BORC subunit sequences using. Small discs indicate individual protein sequences coloured according to subunit assignment (Large discs = colour key).

### BLOC-2 subunits are distantly related to subunits of the Class C tethering complexes HOPS and CORVET

Given the inferred homology between BLOC-1 and BORC, and the previously established relationships between BLOC-3 and the MON1-CCZ1 complex, we next wanted to ascertain whether our broadly representative collection of BLOC-2 subunit sequences could reveal deep relationships to other gene families and detail the origins of this complex.

Surveying the predicted secondary structures and domain architectures of our identified subunits, we found few domains were predicted outside of HPS domains, but approximately 1/3 of HPS5 sequences had C-terminal RING domains, with the distribution broad enough to infer its presence in the LECA (**Figure 3A**, **S4 Table**). The secondary structure of all BLOC-2 subunits comprises an N-terminal region of beta-pleated sheets and a C-terminal region of alpha helices (**Figure 3A**). Within membrane trafficking, this domain organisation is known from subunits of the HOPS, CORVET, SEA, and COPI complexes (Nickerson et al., 2009) (Dokudovskaya and Rout, 2011) (Kaur and Subramanian, 2015). The RING domain has been lost in the HPS5 of both humans and *Drosophila*, while being found in other diverse eukaryotes, including some animals (eg. the brachiopod *Lingula anatine*).

**Figure 3.**
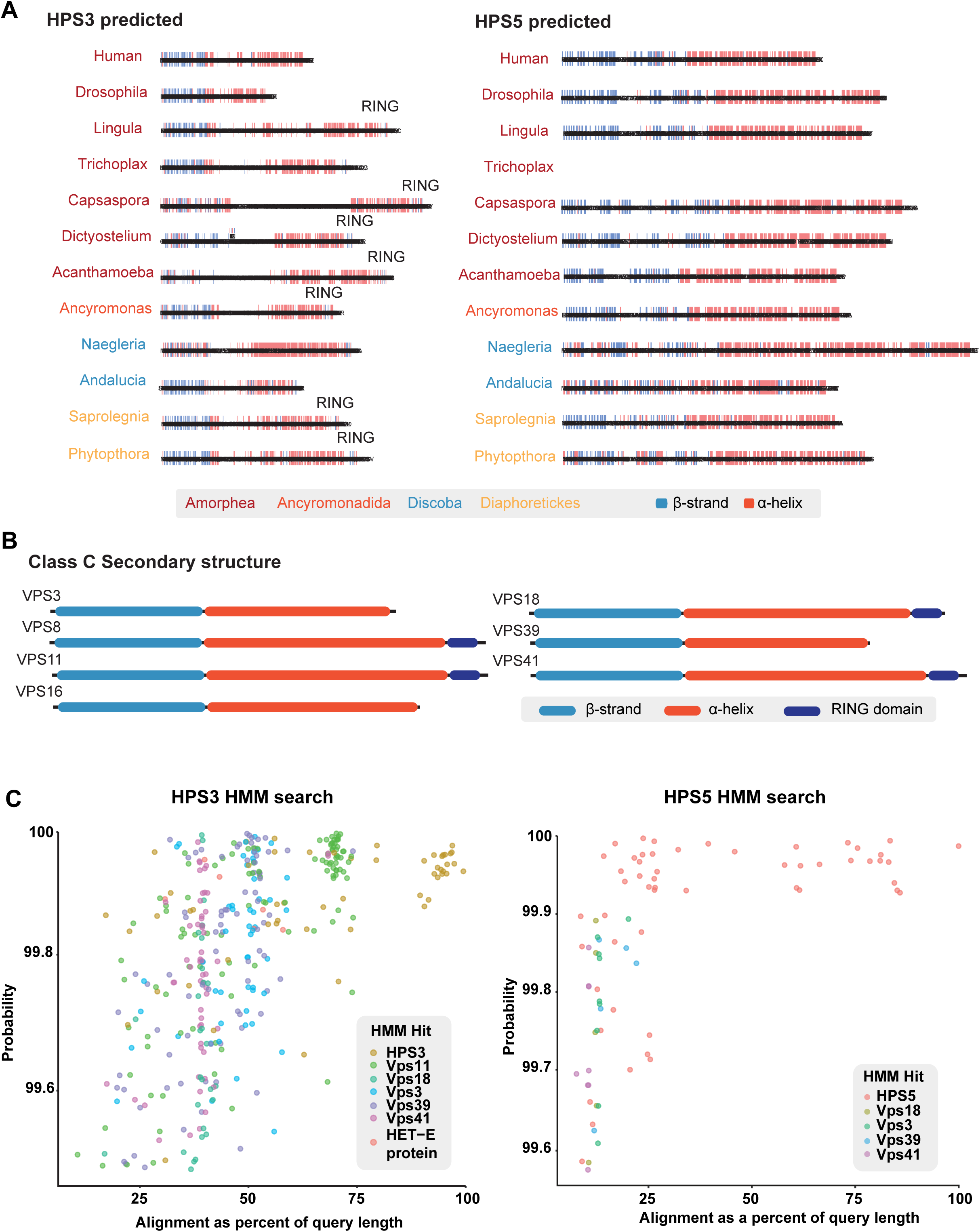
Predicted structural features of BLOC-2 subunits and similarity to Class C tethering factor subunits. **(A)** Secondary structure and RING domain prediction from selected HPS3 and HPS5 subunits of BLOC-2. Full domain prediction data in S4 Table **(B)** Secondary structure and ring domains identified in human HOPS and CORVET Class C tethering factor subunits. **(C)** HMM vs HMM search results using balanced and pan-eukaryotic HPS3 and HPS5 HMMs.

The recently discovered third subunit of the MON1-CCZ1 complex, known as RCM1 in humans and Bulli in *Drosophila* (Vaites et al., 2017) (Dehnen et al., 2020) (van den Boomen et al., 2020) has been speculated to be homologous to HPS6 due to its beta-propeller/alpha solenoid structure (Herrmann et al., 2023). To see if this proposed relationship was supported by sequence similarity, we additionally performed a pan-eukaryotic search for RCM1. Although generally referred to as a metazoan restricted subunit of the complex, RCM1 was found in most genomes along with MON1 and CCZ1, suggesting it is also an ancient and pan-eukaryotic constituent (S3 Table). We used the results to create an RCM1 HMM profile that is more taxonomically diverse than is possible for HPS6 for use in HHblits, a highly sensitive HHM-HMM search tool. We were not able to recover similarity of this subunit to any known proteins; all hits in HHblits were self-hits or unannotated.

BLOC-2 subunits have no detectable sequence similarity to other proteins by BLAST, so we built balanced, eukaryote-wide HMMs to search using HHblits. Results were filtered to exclude hits that solely aligned to the WD40-like N-terminal region. HPS6 retrieved no hits that were not self-hits (S2 Text). HPS3 retrieved most of the HOPS/CORVET subunits (excluding the VpsC proteins Vps16 and Vps8, and the SM protein Vps33), with the strongest similarity to the core subunit Vps11 (**Figure 3C**). Homology between the accessory HOPS/CORVET subunits has previously been proposed (Vps3 and 8 for CORVET, and Vps 39 and 41 for HOPS; Klinger et al 2013); however, our results suggest this homology extends to the accessory subunits and the core subunits Vps11 and Vps18. HPS5 is poorly conserved outside the N-terminal WD40 domain and C-terminal RING domain but retrieved HOPS/CORVET RING domains exclusively (**Figure 3C**). In the case of SEA subunits and alpha-COP, the secondary structure and domain organization was used to propose homology to the HOPS and CORVET subunits (Dokudovskaya and Rout, 2011) (Kaur and Subramanian, 2015); our study is the first to demonstrate direct sequence similarity to HOPS/CORVET in addition to similar secondary structure and domain organization. Our BLOC-2 HMM searches did not retrieve any sequences classified as SEA or alpha-COP, suggesting that we are not simply retrieving proteins of similar secondary structure or domain architecture but are instead identifying deep evolutionary relationships.

## Discussion

Here we have explored the extent and distribution of the LRO-associated complexes from their well-established presence in animals, through the span of eukaryotes and to their deepest evolutionary origins.

Firstly, it is clear that these complexes are far more prevalent and widely spread amongst eukaryotes than currently integrated into cell biological thinking. Our study highlights several protist lineages of agricultural, medical, and economic significance which appear to possess complete or almost complete BLOC complements and may harbour BLOC dependent LROs. Nonetheless, functional data on this pathway beyond the metazoans is currently lacking. Unfortunately, despite there being system-biology resources for some protistan model organisms, none of them possess identifiable BLOC pathways. Moreover, despite using the most sensitive homology-searching tools available, there remains the possibility that some subunits were beyond the limit of bioinformatic detection, but could be identified by direct examination in the organisms of interest, as has been the case for the nuclear pore subunits in *Trypanosoma* (DeGrasse et al., 2009) and more recently COG complex subunits in *Toxoplasma gondii* (Marsilia et al., 2023). With the emerging tools becoming available in an array of protistan model organisms, it should be feasible, fruitful, and exciting to address the nature of BLOC dependent sorting processes in more diverse eukaryotes.

This distribution also strongly implies that all three BLOCs (plus BORC) would have been present in the LECA. Though we identified the BLOCs sparsely, subunits were found in some form in Amorphea + Diaphoretickes (considered on different sides of the eukaryotic root), as well as orphan lineages. Only a few subunits of BLOC-1 and BORC were inferred not to be in the LECA, notably BLOC1S3 (which was specific to Holozoa) and BORCS7 (which was specific to Amorphea and CRuMs). Others, like BLOC1S5, we cannot be confident in due to its extremely sparse distribution. Biochemical characterisation of human BLOC-1 suggests that six subunits of BLOC-1 form two stable, trimeric subcomplexes: the first composed of S1,S4 and S6, and the second of S2, S7 and S8 (Lee et al., 2012). No association of S3 and S5 with either subcomplex was identified, suggesting that these subunits are specifically recruited to the full complex. All subunits in the subcomplexes were inferred to be found in the LECA, suggesting that this subcomplex structure was also present in the LECA, and that BLOC1S3, and perhaps BLOC1S5, were accretions to the functional complex at later stages of eukaryotic evolution. The full BLOC-2 and -3 compliment would have been found in the LECA. This raises the possibility that LECA possessed a bifurcated endosome to lysosome/LRO sorting pathway. Based on our knowledge of the BLOC pathway from animals, this may represent a dedicated regulated secretory pathway in LECA—a major membrane trafficking function that is currently not represented in the pan-eukaryotic “cell plan” of endomembrane organisation. This, however, awaits functional characterization in another eukaryotic model, preferably in a member of the Diaphoretickes, due to their position on the opposite side of the proposed eukaryotic root.

An unanticipated observation was the presence of a C-terminal RING domain in a broad selection of HPS5 proteins. The ancestral HPS5 BLOC-2 subunit likely incorporated a C-terminal RING domain, similar to that found in Class C tether subunits. The RING domains play a role in inter-subunit interactions in the HOPS complex (Hunter et al., 2017) but it does not appear to be essential as this feature has been lost from many yeast HOPS components. The HPS5 RING domain appears to have been lost multiple times throughout eukaryotic evolution as well, including from the model systems in which these complexes are best studied. This underscores the importance of taking a broad comparative approach when attempting to infer the function of the “universal” eukaryotic cell.

Cell biological data suggest that BLOC-2 plays a tethering function (Dennis et al., 2015). However, no sequence relationship to known tethering complexes had previously been identified. We show that BLOC-2 is distantly related to subunits of the Class C tethering complexes, HOPS and CORVET. Whilst HOPS and CORVET are made up of 6 subunits, BLOC-2 consists of only 3 subunits in Amorphea and potentially only two in more distantly related taxa. There is no evidence of BLOC-2 interacting with the Class C core subunits or an SM protein, suggesting a different mechanistic basis for tethering as compared to HOPS and CORVET. However, there is a high degree of plasticity in complex formation between the class-C subunits (van der Beek et al., 2019). Reduced complexes include miniCorvet, identified in *Drosophila,* which lacks the VPS11 core subunit (and the peripheral Vps3, which is lacking from *Drosophila* altogether) (Lőrincz et al., 2016), the CHEVI complex, composed of paralogues of VPS33 and VPS16 in mammals (Cullinane et al., 2010) (Van Der Kant et al., 2015) (Spang, 2016) and a functional dimer of VPS3 and VPS8 involved in membrane traffic between Rab4 positive EE and Rab11 RE in humans (Jonker et al., 2018). Thus, the apparently reduced nature of BLOC-2 when compared to HOPS/CORVET does not preclude its functioning as a homologous tethering complex.

By revealing deep homologies to well characterised post-endosomal trafficking machinery (**Figure 4A**), our work brings the BLOCs—and by extension, some animal LROs— into the organelle paralogy hypothesis in a way that was not possible previously. In addition to the demonstrable sequence similarity and structural similarity between the LRO and endolysosomal pathway components, the functional parallels grow. Most recently is the discovery of BORC’s involvement in the formation of tubules at the phagosome (an endosomal, Rab5 positive organelle similar to the early endosome) that result in trafficking to the lysosome (Fazeli et al., 2023). It was further shown that these BORC-dependent transport carriers fuse with their acceptor membrane through the action of the HOPS tethering complex (Fazeli et al., 2023), mirroring the role of BLOC-1 and BLOC-2 in trafficking from the early endosome to LROs. We propose that this is a bifurcated trafficking pathway from endocytic organelles—BLOC-dependent to LROs, and HOPS/CORVET-dependent to lysosomes—and that these pathways diverged from a primordial endolysosomal trafficking pathway prior to the LECA (**Figure 4B**). A recent analysis of Asgard archaea, the closest prokaryotic relatives of eukaryotes, identified putative HOPS subunits including the core subunits Vps11, 16, and 18, and the accessory subunit Vps39 (Eme et al., 2023). In the event that these are bona-fide Class C subunits, we suggest they are derived from the ancestor of both HOPS/CORVET and BLOC-2 subunits.

**Figure 4.**
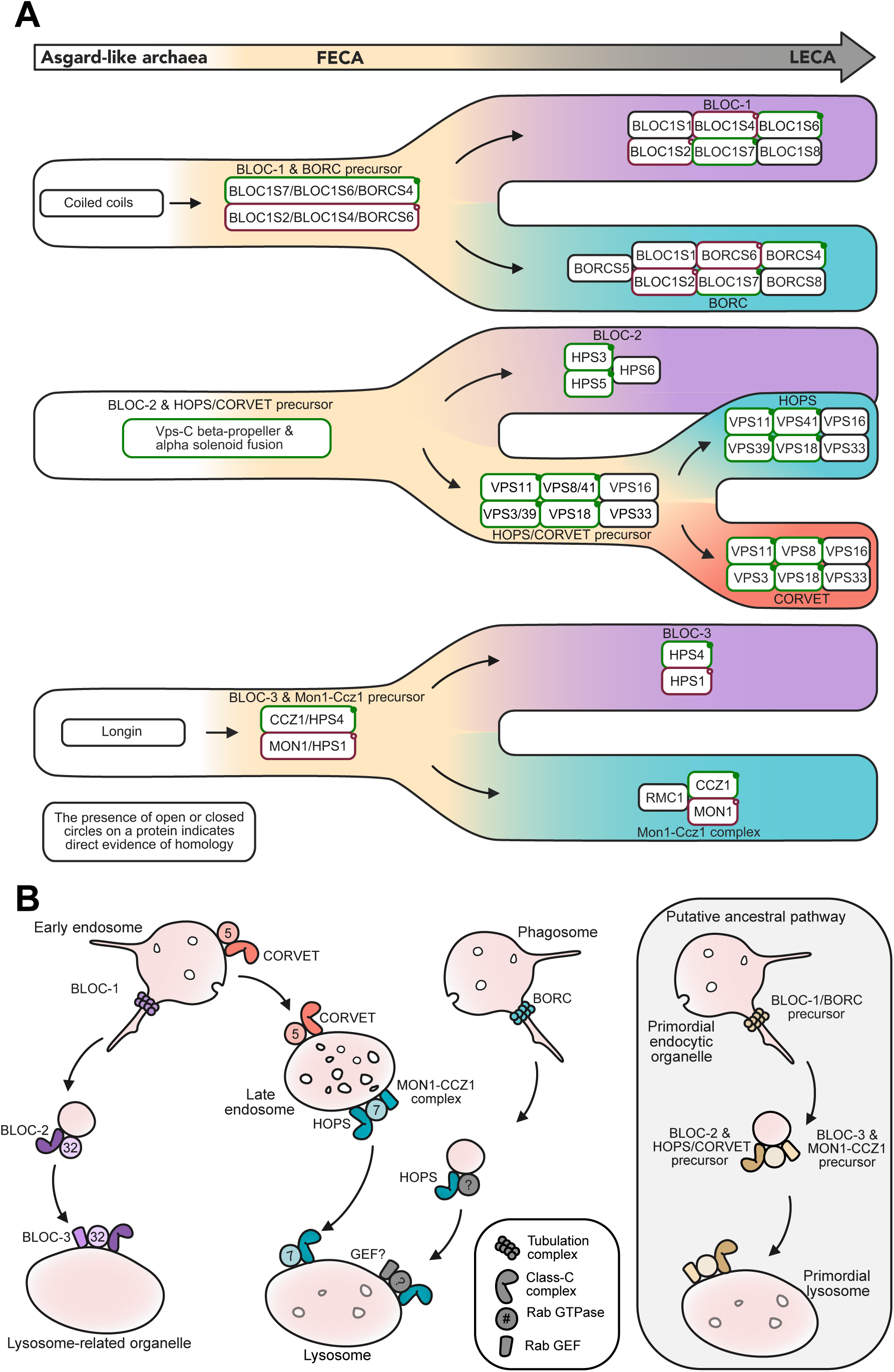
Paralogous complex expansions giving rise to the BLOCs and a model of bifurcation of post-endocytic trafficking during eukaryogenesis. **(A)** Relationships of the BLOCs to other post-endosomal trafficking machinery. Homologous subunit relationships identified in this study and others are denoted by coloured borders. The hypothetical, inferred, deep ancestral precursors are illustrated. The emergence of the full complement of trafficking machinery during the transition from an archaeal ancestor to LECA is shown from left to right. No evolutionary timescale is implied. **(B)** Homologous complexes involved in selected post-endosomal trafficking routes in humans are highlighted. (**Inset**) Our inferred ancestral post-endosomal trafficking pathway, prior to diversification of trafficking routes during later eukaryogenesis, but preceding LECA.

Overall, our work here demonstrates that the LRO-associated trafficking complexes are more widely prevalent and ancient than previously understood. Though their functional roles need to be further explored, we have provided evidence for evolutionary relationships to better characterized post-endosomal trafficking machinery. It is clear that the BLOC dependent sorting pathway needs to be better integrated into our cell biological and evolutionary models of membrane-trafficking in eukaryotes.

## Acknowledgements

We wish to thank B. Franz Lang and Lise Forget for providing the culture of Reclinomonas americana, as well as members of the Dacks Lab and S. Leys for constructive critical input. As well, we thank Sudip Subedi and Jessica Hamilton at the Advanced Cell Exploration Core for technical support and Eleni Karageorgos and Julian Schulz for ongoing administrative support of research in the Dacks Lab.

## Supplementary material

**S1 Dataset. HMMs for genome searches.**

**S1 Text. Transcriptome assembly methods.**

**S2 Text. HPS6 search results.**

**S1 Table. Genome and transcriptome sources used in this study.**

**S2 Table. Full list of positive AMOEBAE search results.**

**S3 Table. Presence/absence distribution of BLOC subunits and related molecular machinery at the species level.** Species represented by genomes are shown in bold, while those represented by transcriptomes are italicised. Positive hits are indicated by ‘+’, except for HPS5 subunits with a predicted RING domain, which are indicated by ‘*’.

**S4 Table. Predicted domains for all HPS5 and TECPR2 positive reciprocal best hits.**

**S1 Figure. Maximum likelihood phylogenies of BLOC-3 subunits and their counterparts from the MON1/CCZ1 complex: HPS1 vs MON1 (A) and HPS4 vs CCZ1 (B).** Support values are 1000 ultrafast bootstraps in IQ-TREE 2. Human sequences in red.

**S2 Figure. Maximum likelihood phylogenies of BLOC-2 subunits HPS3 (A), HPS5 (B), and HPS6 (C). Human sequences in red.**

**S3 Figure. Maximum likelihood phylogeny of BLOC1 subunit BLOC1S6 and core subunit BLOC1S7.** Human sequences in red.

