## Supplemental materials for "The evolutionary origins of the lysosome-related organelle sorting machinery reveal fundamental homology in post-endosome trafficking pathways"

**S1 Text**

***Reclinomonas americana* transcriptome assembly**

Cultures of *Reclinomonas americana* (strain ATCC 50394) were grown in WCL media (https://megasun.bch.umontreal.ca/People/lang/FMGP/methods/wcl.html) at 22 °C and supplied with prey bacterium *Klebsiella aerogenes* (strain ATCC 13048) every 1-3 days. Cultures were harvested using centrifugation, then frozen with liquid nitrogen. RNA was extracted using an Animal Tissue RNA Purification Kit (Norgen Biotek) with an on-column DNase I treatment following the manufacturer's instructions. Libraries were prepared using Illumina TruSeq RNA protocol V2 and sequenced with Illumina Miseq, generating 30 million 75 bp paired end reads.

Reads were trimmed using Bbduk v 38.86 (Bushnell 2020) for quality and adaptor contamination (usejni=t ftm=5 qtrim=rl trimq=5 ktrim=r k=22 mink=11 hdist=1 minlen=35 tpe tbo). Trimmed, paired reads for each sample were then decontaminated with Kraken 2 (Wood and Salzberg 2014; Wood et al. 2019) using the full Kraken 2 standard database from 17/05/2021 (archived at https://benlangmead.github.io/aws-indexes/k2). Reads were error corrected using RCorrector v. 1.0.4 under default settings (Song and Florea 2015), then assembled using rnaSPAdes v 3.15.3 under default settings (Bushmanova et al. 2019). Redundant transcripts were eliminated using cd-hit-est v. 4.8.1 at a global threshold of 98% (Li and Godzik 2006; Fu et al. 2012). The non-redundant set of transcripts was used to predict proteins with GeneMarkS-T v 5.1 (Tang et al., 2015) under default settings, and identical proteins were again collapsed with cd-hit at a global threshold of 100%. Transcriptome quality was assessed with BUSCO v. 5.2.2 in transcriptome mode and eukaryota_odb10 dataset (Simão et al., 2015) (Manni et al., 2021) and TransRate v. 1.0.3 (Smith-Unna et al., 2016) packaged as a part of the Oyster River Protocol Docker

Image (Spillane et al., 2021), https://hub.docker.com/r/macmaneslab/orp and installed as a Singularity image using Singularity version 3.8 (Kurtzer et al., 2017).

**S2 Text**

**HPS6 search results**

The initial BLASTp search of human HPS6 retrieved no positive RBH in the queried genomes. As it is not known to be a human-specific protein and is at least found in other vertebrates (e.g. mice), further investigation was warranted. Human HPS6 was queried into ncbi_nr using BLASTp. A taxonomically diverse selection of the results was then queried against the human genome. Those with a positive reciprocal best hit (RBH) were used to construct a Metazoa-wide HMM (available on Figshare), which was then used to search our selected genomes. This expanded the distribution of HPS6 to at least the other Holozoa (e.g. *Capsaspora owczarzaki*), Apusomonadida (*Thecamonas trahens*) and Amoebozoa (e.g. *Dictyostelium discoideum*) as well.

The distribution of HPS6 was further expanded during an HHblits search that was performed to identify potential deep relations of HPS6 to other protein families. The HPS6 HMM retrieved no annotated profiles that were not HPS6, and closer investigation suggested that many of the unlabelled profiles were unannotated HPS6. These hits were mostly within Amorphea and broadly reflected the distribution of HPS6 sequences found in the AMOEBAE searches described above. A striking exception to this were a handful of profiles for unannotated proteins in oomycetes. RBH was used to identify the orthologs in the genomes used in our study, and when these were aligned and the HMM searched into our annotation database, the sole hit was to the previously identified *Dictyostelium* HPS6. We thus consider these proteins genuine HPS6 proteins.

A

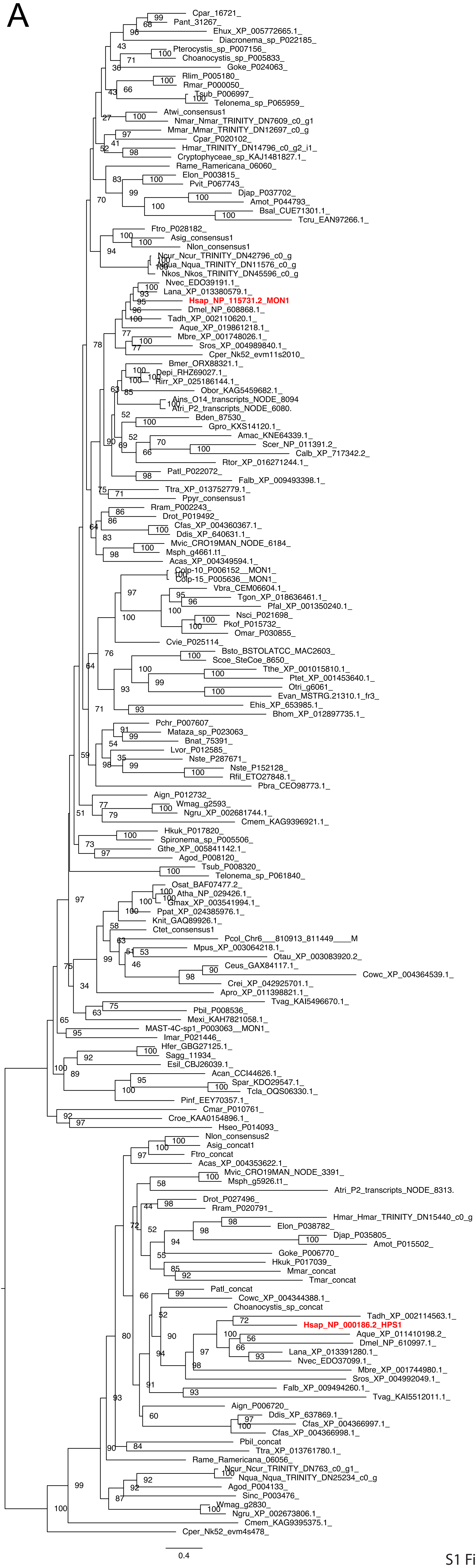
Cpar_16721_
Pant_31267_
Ehux_XP_005772665.1_
Diacronema_sp_P022185_
Pterocystis_sp_P007156_
Choanocystis_sp_P005833_
Goke_P024063_
Rlim_P005180_
Rmar_P000050_
Tsub_P006997_
Telonema_sp_P065959_
Atwi_consensus1
Nmar_Nmar_TRINITY_DN7609_c0_g1
Mmar_Mmar_TRINITY_DN12697_c0_g
Cpar_P020102_
Hmar_TRINITY_DN14796_c0_g2_i1_
Cryptophyceae_sp_KAJ1481827.1_
Rame_Ramericana_06060_
Elon_P003815_
Pvit_P067743_
Djap_P037702_
Amot_P044793_
Bsal_CUE71301.1_
Tcru_EAN97266.1_
Ftro_P028182_
Asig_consensus1
Nlon_consensus1
Ncur_Ncur_TRINITY_DN42796_c0_g
Nqua_Nqua_TRINITY_DN11576_c0_g
Nkos_Nkos_TRINITY_DN45596_c0_g
Nvec_EDO39191.1_
Lana_XP_013380579.1_
Hsap_NP_115731.2_MON1
Dmel_NP_608868.1_
Tadh_XP_002110620.1_
Aque_XP_019861218.1_
Mbre_XP_001748026.1_
Sros_XP_004989840.1_
Cper_Nk52_evm11s2010_
Bmer_ORX88321.1_
Depi_RHZ69027.1_
Rirr_XP_025186144.1_
Obor_KAG5459682.1_
Ains_O14_transcripts_NODE_8094
Atri_P2_transcripts_NODE_6080.
Bden_87530_
Gpro_KXS14120.1_
Amac_KNE64339.1_
Scer_NP_011391.2_
Calb_XP_717342.2_
Rtor_XP_016271244.1_
Patl_P022072_
Falb_XP_009493398.1_
Ttra_XP_013752779.1_
Ppyr_consensus1
Rram_P002243_
Drot_P019492_
Cfas_XP_004360367.1_
Ddis_XP_640631.1_
Mvic_CRO19MAN_NODE_6184_
Msph_g4661.t1_
Acas_XP_004349594.1_
Colp-10_P006152__MON1_
Colp-15_P005636__MON1_
Vbra_CEM06604.1_
Tgon_XP_018636461.1_
Pfal_XP_001350240.1_
Nsci_P021698_
Pkof_P015732_
Omar_P030855_
Cvie_P025114_
Bsto_BSTOLATCC_MAC2603_
Scoe_SteCoe_8650_
Tthe_XP_001015810.1_
Ptet_XP_001453640.1_
Otri_g6061_
Evan_MSTRG.21310.1_fr3_
Ehis_XP_653985.1_
Bhom_XP_012897735.1_
Pchr_P007607_
Mataza_sp_P023063_
Bnat_75391_
Lvor_P012585_
Nste_P287671_
Nste_P152128_
Rfil_ETO27848.1_
Pbra_CEO98773.1_
Aign_P012732_
Wmag_g2593_
Ngru_XP_002681744.1_
Cmem_KAG9396921.1_
Hkuk_P017820_
Spironema_sp_P005506_
Gthe_XP_005841142.1_
Agod_P008120_
Tsub_P008320_
Telonema_sp_P061840_
Osat_BAF07477.2_
Atha_NP_029426.1_
Gmax_XP_003541994.1_
Ppat_XP_024385976.1_
Knit_GAQ89926.1_
Ctet_consensus1
Pcol_Chr6___810913_811449____M
Mpus_XP_003064218.1_
Otau_XP_003083920.2_
Ceus_GAX84117.1_
Cowc_XP_004364539.1_
Crei_XP_042925701.1_
Apro_XP_011398821.1_
Tvag_KAI5496670.1_
Pbil_P008536_
Mexi_KAH7821058.1_
MAST-4C-sp1_P003063__MON1_
Imar_P021446_
Hfer_GBG27125.1_
Sagg_11934_
Esil_CBJ26039.1_
Acan_CCI44626.1_
Spar_KDO29547.1_
Tcla_OQS06330.1_
Pinf_EEY70357.1_
Cmar_P010761_
Croe_KAA0154896.1_
Hseo_P014093_
Nlon_consensus2
Asig_concat1
Ftro_concat
Acas_XP_004353622.1_
Mvic_CRO19MAN_NODE_3391_
Msph_g5926.t1_
Atri_P2_transcripts_NODE_8313.
Drot_P027496_
Rram_P020791_
Hmar_Hmar_TRINITY_DN15440_c0_g
Elon_P038782_
Djap_P035805_
Amot_P015502_
Goke_P006770_
Hkuk_P017039_
Mmar_concat
Tmar_concat
Patl_concat
Cowc_XP_004344388.1_
Choanocystis_sp_concat
Tadh_XP_002114563.1_
Hsap_NP_000186.2_HPS1
Aque_XP_011410198.2_
Dmel_NP_610997.1_
Lana_XP_013391280.1_
Nvec_EDO37099.1_
Mbre_XP_001744980.1_
Sros_XP_004992049.1_
Falb_XP_009494260.1_
Tvag_KAI5512011.1_
Aign_P006720_
Ddis_XP_637869.1_
Cfas_XP_004366997.1_
Cfas_XP_004366998.1_
Pbil_concat
Ttra_XP_013761780.1_
Rame_Ramericana_06056_
Ncur_Ncur_TRINITY_DN763_c0_g1_
Nqua_Nqua_TRINITY_DN25234_c0_g
Agod_P004133_
Sinc_P003476_
Wmag_g2830_
Ngru_XP_002673806.1_
Cmem_KAG9395375.1_
Cper_Nk52_evm4s478_
0.4

S1 Figure

B

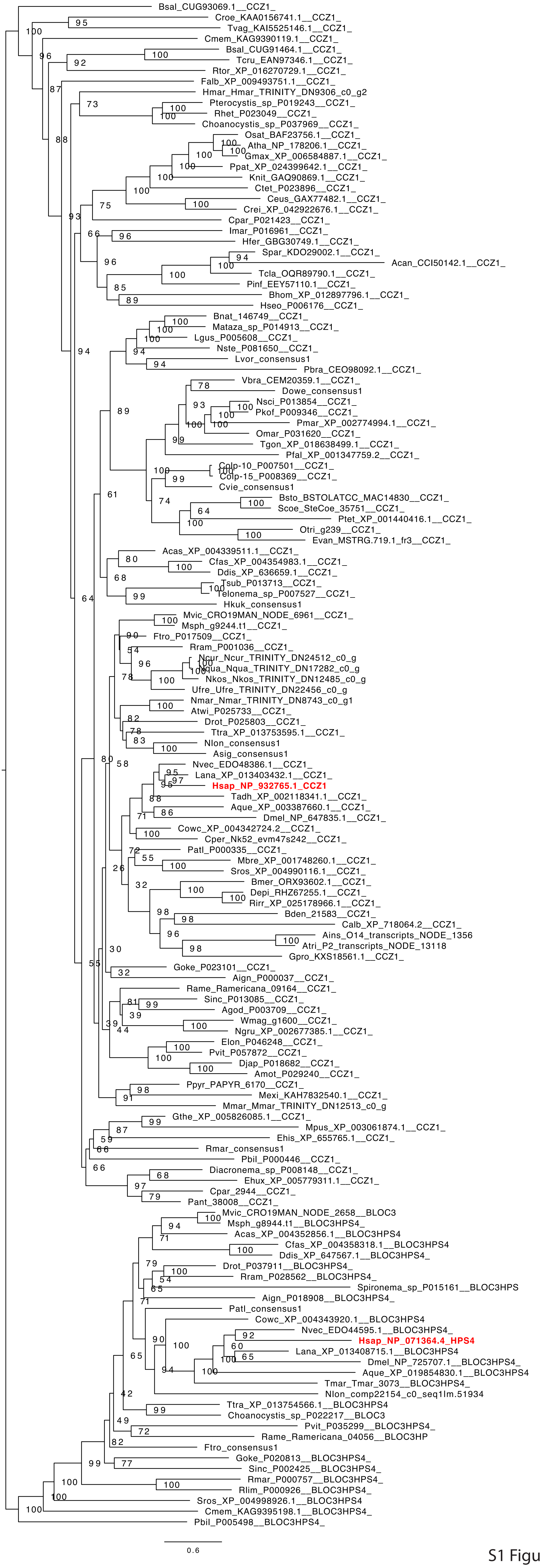

Bsal_CUG93069.1__CCZ1_
Croe_KAA0156741.1__CCZ1_
Tvag_KAI5525146.1__CCZ1_
Cmem_KAG9390119.1__CCZ1_
Bsal_CUG91464.1__CCZ1_
Tcru_EAN97346.1__CCZ1_
Rtor_XP_016270729.1__CCZ1_
Falb_XP_009493751.1__CCZ1_
Hmar_Hmar_TRINITY_DN9306_c0_g2
Pterocystis_sp_P019243__CCZ1_
Rhet_P023049__CCZ1_
Choanocystis_sp_P037969__CCZ1_
Osat_BAF23756.1__CCZ1_
Atha_NP_178206.1__CCZ1_
Gmax_XP_006584887.1__CCZ1_
Ppat_XP_024399642.1__CCZ1_
Knit_GAQ90869.1__CCZ1_
Ctet_P023896__CCZ1_
Ceus_GAX77482.1__CCZ1_
Crei_XP_042922676.1__CCZ1_
Cpar_P021423__CCZ1_
Imar_P016961__CCZ1_
Hfer_GBG30749.1__CCZ1_
Spar_KDO29002.1__CCZ1_
Acan_CCI50142.1__CCZ1_
Tcla_OQR89790.1__CCZ1_
Pinf_EEY57110.1__CCZ1_
Bhom_XP_012897796.1__CCZ1_
Hseo_P006176__CCZ1_
Bnat_146749__CCZ1_
Mataza_sp_P014913__CCZ1_
Lgus_P005608__CCZ1_
Nste_P081650__CCZ1_
Lvor_consensus1
Pbra_CEO98092.1__CCZ1_
Vbra_CEM20359.1__CCZ1_
Dowe_consensus1
Nsci_P013854__CCZ1_
Pkof_P009346__CCZ1_
Pmar_XP_002774994.1__CCZ1_
Omar_P031620__CCZ1_
Tgon_XP_018638499.1__CCZ1_
Pfal_XP_001347759.2__CCZ1_
Colp-10_P007501__CCZ1_
Colp-15_P008369__CCZ1_
Cvie_consensus1
Bsto_BSTOLATCC_MAC14830__CCZ1_
Scoe_SteCoe_35751__CCZ1_
Ptet_XP_001440416.1__CCZ1_
Otri_g239__CCZ1_
Evan_MSTRG.719.1_fr3__CCZ1_
Acas_XP_004339511.1__CCZ1_
Cfas_XP_004354983.1__CCZ1_
Ddis_XP_636659.1__CCZ1_
Tsub_P013713__CCZ1_
Telonema_sp_P007527__CCZ1_
Hkuk_consensus1
Mvic_CRO19MAN_NODE_6961__CCZ1_
Msph_g9244.t1__CCZ1_
Ftro_P017509__CCZ1_
Rram_P001036__CCZ1_
Ncur_Ncur_TRINITY_DN24512_c0_g
Nqua_Nqua_TRINITY_DN17282_c0_g
Nkos_Nkos_TRINITY_DN12485_c0_g
Ufre_Ufre_TRINITY_DN22456_c0_g
Nmar_Nmar_TRINITY_DN8743_c0_g1
Atwi_P025733__CCZ1_
Drot_P025803__CCZ1_
Ttra_XP_013753595.1__CCZ1_
Nlon_consensus1
Asig_consensus1
Nvec_EDO48386.1__CCZ1_
Lana_XP_013403432.1__CCZ1_
Hsap_NP_932765.1_CCZ1
Tadh_XP_002118341.1__CCZ1_
Aque_XP_003387660.1__CCZ1_
Dmel_NP_647835.1__CCZ1_
Cowc_XP_004342724.2__CCZ1_
Cper_Nk52_evm47s242__CCZ1_
Patl_P000335__CCZ1_
Mbre_XP_001748260.1__CCZ1_
Sros_XP_004990116.1__CCZ1_
Bmer_ORX93602.1__CCZ1_
Depi_RHZ67255.1__CCZ1_
Rirr_XP_025178966.1__CCZ1_
Bden_21583__CCZ1_
Calb_XP_718064.2__CCZ1_
Ains_O14_transcripts_NODE_1356
Atri_P2_transcripts_NODE_13118
Gpro_KXS18561.1__CCZ1_
Goke_P023101__CCZ1_
Aign_P000037__CCZ1_
Rame_Ramericana_09164__CCZ1_
Sinc_P013085__CCZ1_
Agod_P003709__CCZ1_
Wmag_g1600__CCZ1_
Ngru_XP_002677385.1__CCZ1_
Elon_P046248__CCZ1_
Pvit_P057872__CCZ1_
Djap_P018682__CCZ1_
Amot_P029240__CCZ1_
Ppyr_PAPYR_6170__CCZ1_
Mexi_KAH7832540.1__CCZ1_
Mmar_Mmar_TRINITY_DN12513_c0_g
Gthe_XP_005826085.1__CCZ1_
Mpus_XP_003061874.1__CCZ1_
Ehis_XP_655765.1__CCZ1_
Rmar_consensus1
Pbil_P000446__CCZ1_
Diacronema_sp_P008148__CCZ1_
Ehux_XP_005779311.1__CCZ1_
Cpar_2944__CCZ1_
Pant_38008__CCZ1_
Mvic_CRO19MAN_NODE_2658__BLOC3
Msph_g8944.t1__BLOC3HPS4_
Acas_XP_004352856.1__BLOC3HPS4
Cfas_XP_004358318.1__BLOC3HPS4
Ddis_XP_647567.1__BLOC3HPS4_
Drot_P037911__BLOC3HPS4_
Rram_P028562__BLOC3HPS4_
Spironema_sp_P015161__BLOC3HPS
Aign_P018908__BLOC3HPS4_
Patl_consensus1
Cowc_XP_004343920.1__BLOC3HPS4
Nvec_EDO44595.1__BLOC3HPS4_
Hsap_NP_071364.4_HPS4
Lana_XP_013408715.1__BLOC3HPS4
Dmel_NP_725707.1__BLOC3HPS4_
Aque_XP_019854830.1__BLOC3HPS4
Tmar_Tmar_3073__BLOC3HPS4_
Nlon_comp22154_c0_seq1lm.51934
Ttra_XP_013754566.1__BLOC3HPS4
Choanocystis_sp_P022217__BLOC3
Pvit_P035299__BLOC3HPS4_
Rame_Ramericana_04056__BLOC3HP
Ftro_consensus1
Goke_P020813__BLOC3HPS4_
Sinc_P002425__BLOC3HPS4_
Rmar_P000757__BLOC3HPS4_
Rlim_P000926__BLOC3HPS4_
Sros_XP_004998926.1__BLOC3HPS4
Cmem_KAG9395198.1__BLOC3HPS4_
Pbil_P005498__BLOC3HPS4_
0.6


S1 Figure

A

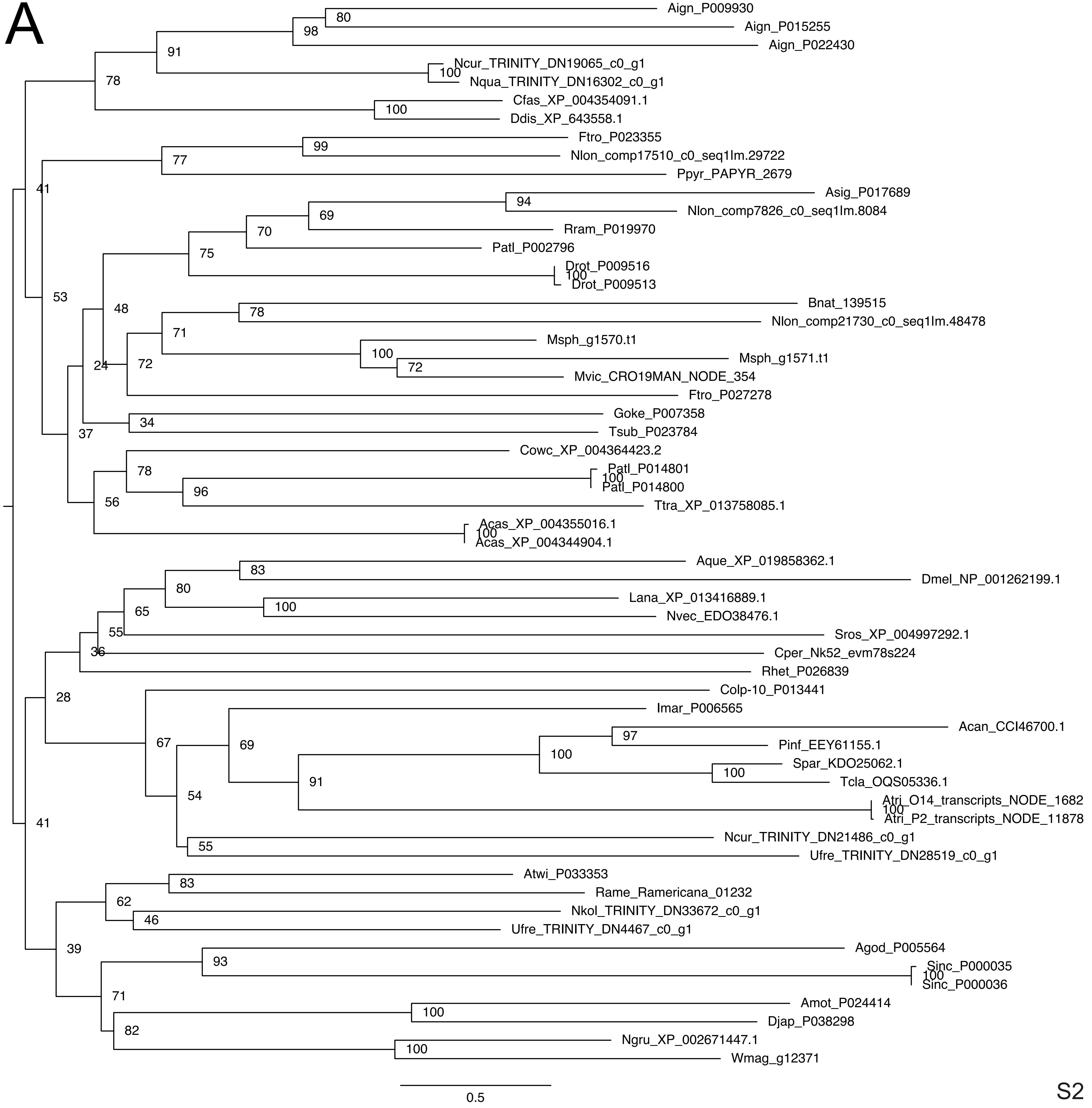
Aign_P009930
Aign_P015255
Aign_P022430
Ncur_TRINITY_DN19065_c0_g1
Nqua_TRINITY_DN16302_c0_g1
Cfas_XP_004354091.1
Ddis_XP_643558.1
Ftro_P023355
Nlon_comp17510_c0_seq1lm.29722
Ppyr_PAPYR_2679
Asig_P017689
Nlon_comp7826_c0_seq1lm.8084
Rram_P019970
Patl_P002796
Drot_P009516
Drot_P009513
Bnat_139515
Nlon_comp21730_c0_seq1lm.48478
Msph_g1570.t1
Msph_g1571.t1
Mvic_CRO19MAN_NODE_354
Ftro_P027278
Goke_P007358
Tsub_P023784
Cowc_XP_004364423.2
Patl_P014801
Patl_P014800
Ttra_XP_013758085.1
Acas_XP_004355016.1
Acas_XP_004344904.1
Aque_XP_019858362.1
Dmel_NP_001262199.1
Lana_XP_013416889.1
Nvec_EDO38476.1
Sros_XP_004997292.1
Cper_Nk52_evm78s224
Rhet_P026839
Colp-10_P013441
Imar_P006565
Acan_CCI46700.1
Pinf_EEY61155.1
Spar_KDO25062.1
Tcla_OQS05336.1
Atri_O14_transcripts_NODE_1682
Atri_P2_transcripts_NODE_11878
Ncur_TRINITY_DN21486_c0_g1
Ufre_TRINITY_DN28519_c0_g1
Atwi_P033353
Rame_Ramericana_01232
Nkol_TRINITY_DN33672_c0_g1
Ufre_TRINITY_DN4467_c0_g1
Agod_P005564
Sinc_P000035
Sinc_P000036
Amot_P024414
Djap_P038298
Ngru_XP_002671447.1
Wmag_g12371
0.5

S2 Figure

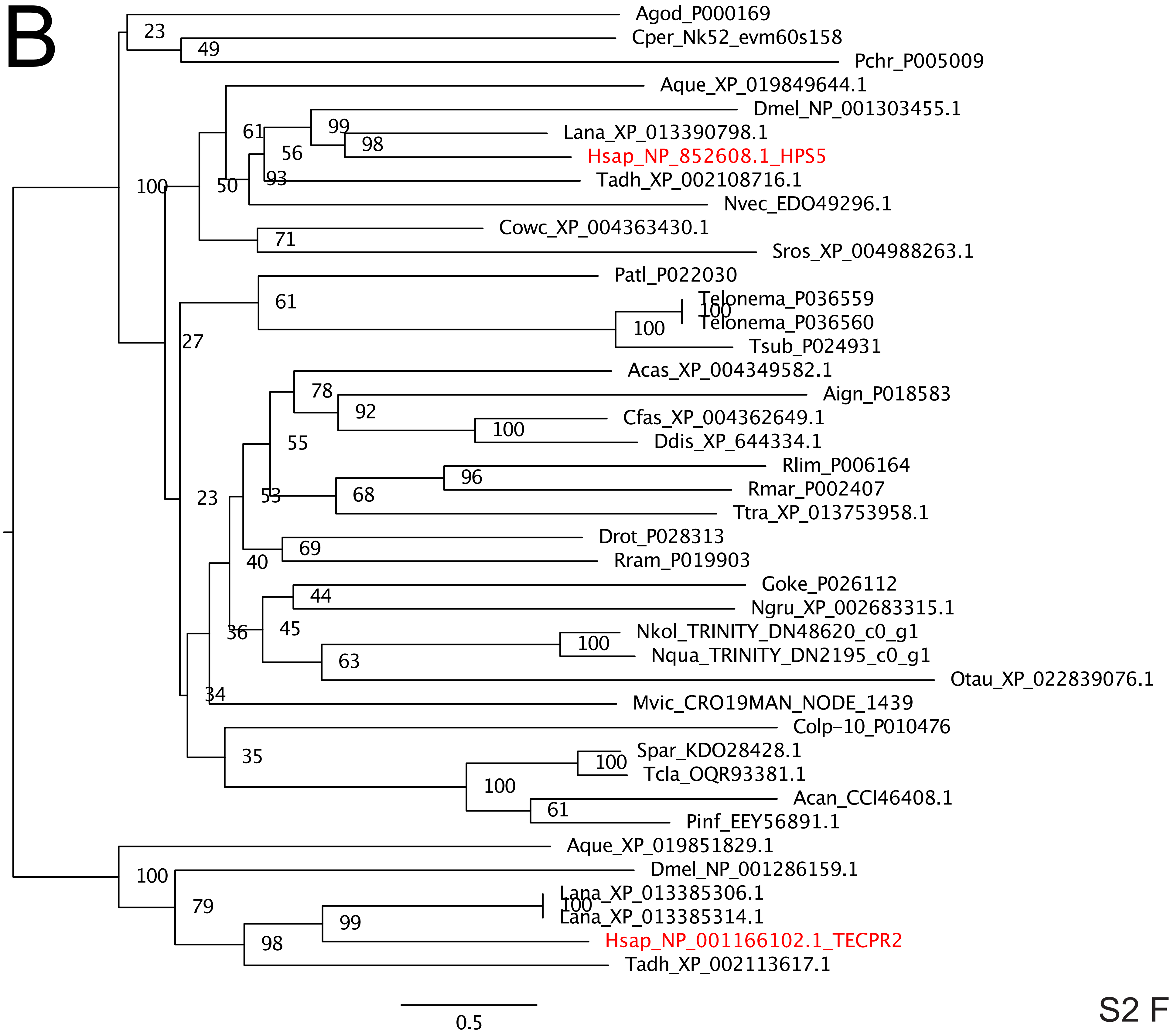
B
Agod_P000169
Cper_Nk52_evm60s158
Pchr_P005009
Aque_XP_019849644.1
Dmel_NP_001303455.1
Lana_XP_013390798.1
Hsap_NP_852608.1_HPS5
Tadh_XP_002108716.1
Nvec_EDO49296.1
Cowc_XP_004363430.1
Sros_XP_004988263.1
Patl_P022030
Telonema_P036559
Telonema_P036560
Tsub_P024931
Acas_XP_004349582.1
Aign_P018583
Cfas_XP_004362649.1
Ddis_XP_644334.1
Rlim_P006164
Rmar_P002407
Ttra_XP_013753958.1
Drot_P028313
Rram_P019903
Goke_P026112
Ngru_XP_002683315.1
Nkol_TRINITY_DN48620_c0_g1
Nqua_TRINITY_DN2195_c0_g1
Otau_XP_022839076.1
Mvic_CRO19MAN_NODE_1439
Colp-10_P010476
Spar_KDO28428.1
Tcla_OQR93381.1
Acan_CCI46408.1
Pinf_EEY56891.1
Aque_XP_019851829.1
Dmel_NP_001286159.1
Lana_XP_013385306.1
Lana_XP_013385314.1
Hsap_NP_001166102.1_TECPR2
Tadh_XP_002113617.1
0.5

S2 Figure

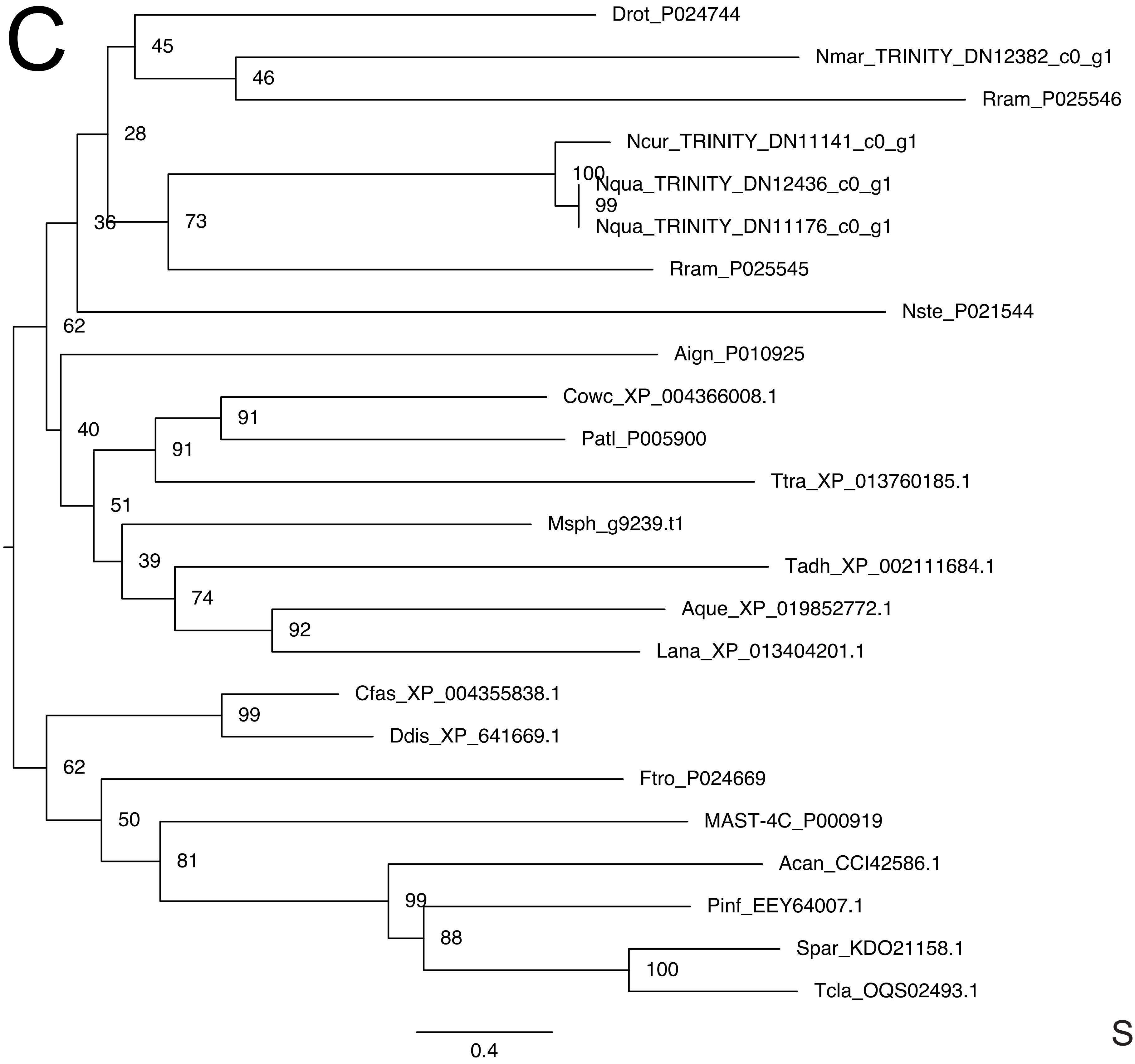
C
Drot_P024744
Nmar_TRINITY_DN12382_c0_g1
Rram_P025546
Ncur_TRINITY_DN11141_c0_g1
Nqua_TRINITY_DN12436_c0_g1
Nqua_TRINITY_DN11176_c0_g1
Rram_P025545
Nste_P021544
Aign_P010925
Cowc_XP_004366008.1
Patl_P005900
Ttra_XP_013760185.1
Msph_g9239.t1
Tadh_XP_002111684.1
Aque_XP_019852772.1
Lana_XP_013404201.1
Cfas_XP_004355838.1
Ddis_XP_641669.1
Ftro_P024669
MAST-4C_P000919
Acan_CCI42586.1
Pinf_EEY64007.1
Spar_KDO21158.1
Tcla_OQS02493.1
45
46
28
100
99
36
73
62
40
91
91
51
39
74
92
99
62
50
81
99
88
100
0.4

S2 Figure

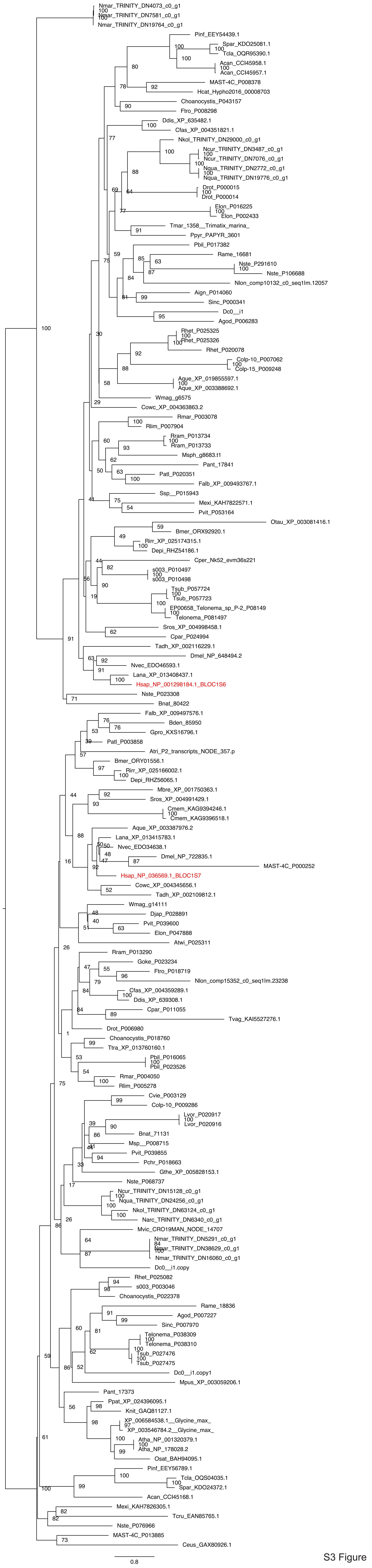

S3 Figure
